# Targeted deletion of Pf prophages from diverse *Pseudomonas aeruginosa* isolates impacts quorum sensing and virulence traits

**DOI:** 10.1101/2023.11.19.567716

**Authors:** Amelia K. Schmidt, Caleb M. Schwartzkopf, Julie D. Pourtois, Elizabeth Burgener, Dominick R. Faith, Alex Joyce, Tyrza Lamma, Geetha Kumar, Paul L. Bollyky, Patrick R. Secor

## Abstract

*Pseudomonas aeruginosa* is an opportunistic bacterial pathogen that commonly causes medical hardware, wound, and respiratory infections. Temperate filamentous Pf phages that infect *P. aeruginosa* impact numerous bacterial virulence phenotypes. Most work on Pf phages has focused on strain Pf4 and its host *P. aeruginosa* PAO1. Expanding from Pf4 and PAO1, this study explores diverse Pf strains infecting *P. aeruginosa* clinical isolates. We describe a simple technique targeting the Pf lysogeny maintenance gene, *pflM* (*PA0718*), that enables the effective elimination of Pf prophages from diverse *P. aeruginosa* hosts. This study also assesses the effects different Pf phages have on host quorum sensing, biofilm formation, virulence factor production, and virulence. Collectively, this research not only introduces a valuable tool for Pf prophage elimination from diverse *P. aeruginosa* isolates, but also advances our understanding of the complex relationship between *P. aeruginosa* and filamentous Pf phages.

**Importance:** *Pseudomonas aeruginosa* is an opportunistic bacterial pathogen that is frequently infected by filamentous Pf phages (viruses) that integrate into its chromosome, affecting behavior. While prior work has focused on Pf4 and PAO1, this study investigates diverse Pf strains in clinical isolates. A simple method targeting the deletion of the Pf lysogeny maintenance gene *pflM* (*PA0718*) effectively eliminates Pf prophages from clinical isolates. The research evaluates the impact Pf prophages have on bacterial quorum sensing, biofilm formation, and virulence phenotypes. This work introduces a valuable tool to eliminate Pf prophages from clinical isolates and advances our understanding of *P. aeruginosa* and filamentous Pf phage interactions.

## Introduction

*Pseudomonas aeruginosa* is an opportunistic bacterial pathogen that commonly infects medical hardware, diabetic ulcers, burn wounds, and the airways of cystic fibrosis patients (1). *P. aeruginosa* isolates are often infected by filamentous viruses (phages) called Pf (2-4). Pf phages live a temperate lifestyle and integrate into the bacterial chromosome as a prophage, passively replicating with each bacterial cell division. When induced, the Pf prophage is excised from the chromosome forming a circular double-stranded episome called the replicative form (5). Pf replicative form copy numbers increase in the cytoplasm where they serve as templates for viral transcription and the production of circular single-stranded DNA genomes that are packaged into filamentous virions as they are extruded from the cell by a process that is analogous to type IV pili assembly (3, 6).

Filamentous Pf virions enhance *P. aeruginosa* virulence potential by promoting biofilm formation (7) and inhibiting phagocytic uptake by macrophages (8, 9). Pf virions also carry a high negative charge density allowing them to sequester cationic antimicrobials such as aminoglycoside antibiotics and antimicrobial peptides (7, 10, 11). Additionally, Pf phages enhance the virulence potential of *P. aeruginosa* by modulating the secretion of the quorum-regulated virulence factor pyocyanin (12, 13). These properties may explain why the presence of Pf virions at sites of infection is associated with more chronic lung infections and antibiotic resistance in cystic fibrosis patients (8) and why *P. aeruginosa* strains cured of their Pf infection are less virulent in murine models of pneumonia (14) and wound infection (15).

Most studies to date have focused on interactions between Pf strain Pf4 and its host *P. aeruginosa* PAO1. (3, 9, 11, 14). Despite the clear link between Pf4 and the virulence of *P. aeruginosa* PAO1, the effects diverse Pf strains that infect *P. aeruginosa* clinical isolates have on virulence phenotypes remains unclear. This is in part due to the significant challenge of ‘curing’ clinical isolates of their Pf prophage infections. Prior efforts to delete Pf4 from PAO1 relied on the integration of a selectable marker into the integration site used by Pf4 (14), which precludes complementation studies that re-introduce the Pf4 prophage to the host chromosome. In prior work, we were able to generate a clean Pf4 deletion strain by first deleting the *pfiTA* toxin-antitoxin module encoded by Pf4 followed by deletion of the rest of the prophage (16).

Here, we find that the Pf4 gene *PA0718* maintains Pf4 in a lysogenic state; we therefore refer to *PA0718* as the <u>Pf l</u>ysogeny <u>m</u>aintenance gene *pflM*. Deletion of *PA0718* or homologous alleles from Pf prophages in clinical *P. aeruginosa* isolates LESB58, CPA0053, CPA0087, and the multidrug resistant strain DDRC3 resulted in the complete loss of Pf prophages from each strain. Furthermore, we observe that some substrains of PAO1 are lysogenized by two Pf phages, Pf4 and Pf6, and we successfully cured PAO1 of both Pf4 and Pf6 prophages. We compare phenotypic differences between wild-type and ΔPf prophage mutants by assessing Las, Rhl, and PQS quorum sensing activity, biofilm formation, and pyocyanin production. We also examine how Pf prophages impact virulence phenotypes in a *Caenorhabditis elegans* avoidance model. Overall, we present a new methodology for efficiently curing *P. aeruginosa* strains of their resident Pf prophages and leverage this tool to gain insight into the diverse impacts Pf phages have on their bacterial hosts.

## Results

### PA0718 (PflM) maintains Pf4 integration in *P. aeruginosa* PAO1

While making single gene deletions from the core Pf4 genome (*pf4r-intF4)* in *P. aeruginosa* PAO1 (**Fig. 1A**), we noted that deleting either the *Pf4r* repressor or the *PA0718* gene results in the complete excision of the Pf4 prophage from the *P. aeruginosa* chromosome (**Fig. 1B**, upper bands). Prior work demonstrates that deletion of the *Pf4r* repressor induces Pf4 prophage excision and virion replication (17), but how *PA0718* is involved in Pf4 excision is not known.

**Figure 1:**
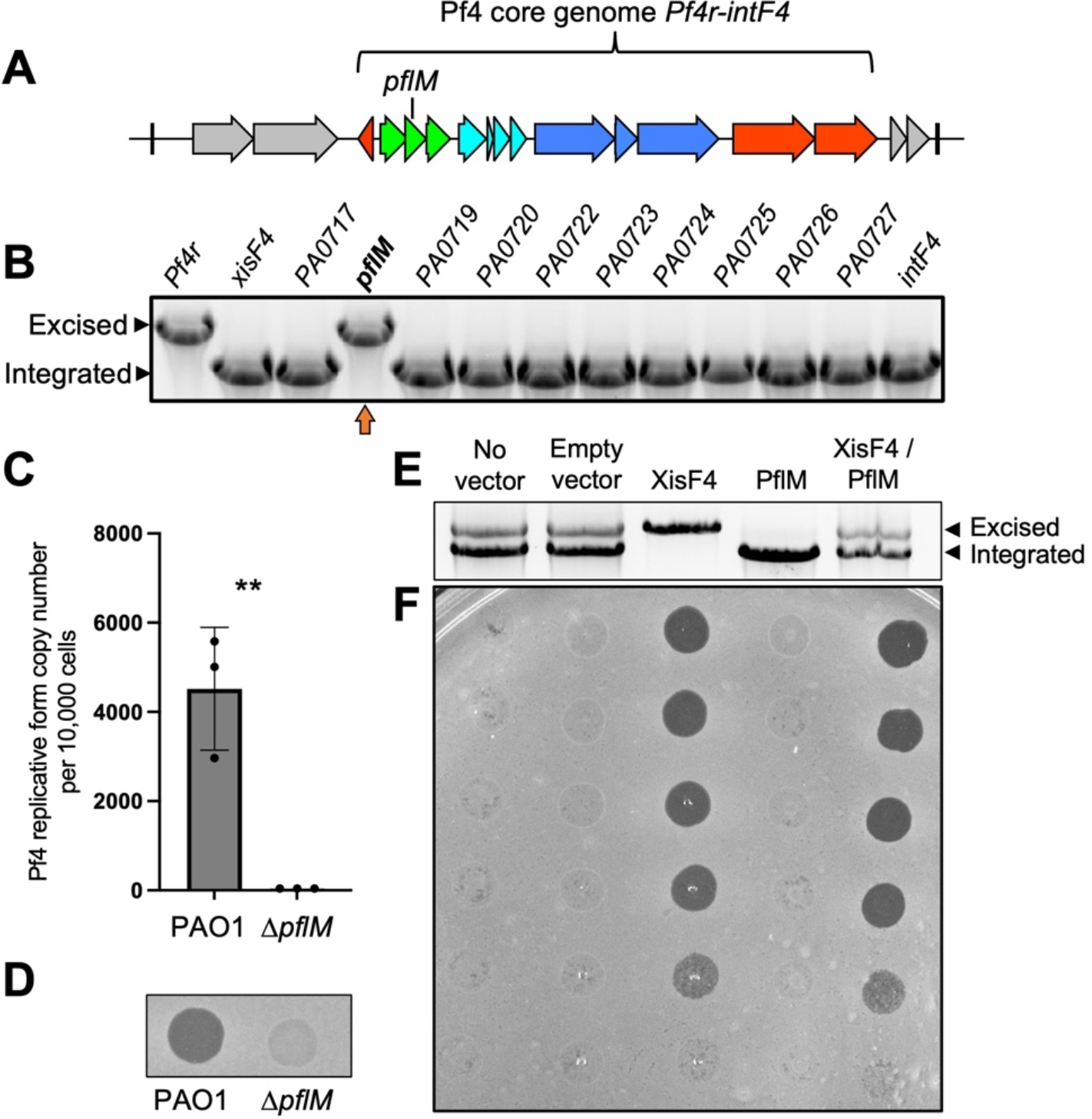
PA0718 (PflM) maintains Pf4 lysogeny. **(A)** Schematic of the Pf4 prophage (*PA0715-PA0729*) integrated into the tRNA-Gly gene *PA0729*.*1* in *P. aeruginosa* PAO1. The core genome that is conserved amongst Pf strains is indicated. **(B)** Multiplex PCR was used to measure Pf4 prophage integration and excision from the PAO1 chromosome in the indicated Pf4 single-gene mutants. Excision and integration are differentiated by the size of the PCR product produced. Note that that deleting *PA0721* (*pfsE*) from the Pf4 prophage is lethal to *P. aeruginosa* (16) explaining why *pfsE* is not included in the assay. A representative gel is shown. **(C)** Quantitative PCR (qPCR) was used to measure episomal Pf4 replicative form in PAO1 or Δ*PA0718* cells after 18 hours of growth in LB broth. Data are the mean of three replicate experiments, Δ*PA0718* values were below the limit of detection for the assay (37 copies per microliter of input material). **(D)** Supernatants obtained from 18 hour-old cultures of PAO1 or Δ*PA0718* were spotted onto lawns of *P. aeruginosa* ΔPf4. A representative image is shown. **(E)** PflM and/or XisF4 were expressed from inducible plasmids in *P. aeruginosa* PAO1. After 18 hours of growth in LB broth, Pf4 integration and excision was measured by multiplex PCR. **(F)** Filtered supernatants collected from the indicated strains were tittered on lawns of PAO1^ΔPf4^ and imaged after 18 hours of growth.

After excision, Pf4 replicates as a circular episome called the replicative form (5). We used qPCR to measure circular Pf4 replicative form copy number in wild-type and Δ*PA0718* cells. In wild-type cells, approximately 4,400 replicative form copies were detected for every 10,000 cells; however, the Pf4 replicative form was not detected in Δ*PA0718* cells (**Fig. 1C**), indicating that Pf4 genome replication is not initiated, and the replicative form is lost as cells divide. Consistently, infectious Pf4 virions are detected in supernatants collected from wild-type cultures but not in supernatants collected from Δ*PA0718* (**Fig. 1D**). These results indicate that PA0718 maintains the Pf4 prophage in a lysogenic state and that deleting *PA0718* induces Pf4 prophage excision, but not replication, curing PAO1 of its Pf4 infection. Herein, we refer to *PA0718* as the <u>Pf</u> lysogeny <u>m</u>aintenance gene *pflM*.

The observation that 4,400 Pf4 replicative form copies are detected for every 10,000 wild-type cells (**Fig. 1C**) indicates Pf4 is actively replicating in a subpopulation of cells. We used a multiplex PCR excision assay to measure Pf4 prophage excision and integration in *P. aeruginosa* populations. In PAO1 populations with no expression vector or those carrying an empty expression vector, both Pf4 prophage integration and excision are observed (**Fig. 1E**, two bands are present); however, infectious virions were not detected in supernatants by plaque assay (**Fig. 1F**), suggesting that Pf4 is replicating at low levels during planktonic growth in LB broth, consistent with prior results (18).

The Pf4 excisionase XisF4 regulates Pf4 prophage excision (17) and expressing XisF4 in *trans* induces complete Pf4 prophage excision (**Fig 1E**) and robust virion replication (**Fig. 1F**). In contrast, expressing PflM in *trans* maintains the entire population in a lysogenic state (**Fig. 1E**) and virion replication is not detected (**Fig. 1F**). When PflM and XisF4 are expressed together, both Pf4 integration and excision products are observed (**Fig. 1E**) and infectious virions are produced at titers comparable to cells where XisF4 was expressed by itself (**Fig. 1F**). These results indicate that expressing PflM is not sufficient to inhibit XisF4-mediated Pf4 prophage excision and replication, but that PflM can maintain some cells in a lysogenic state during active viral replication.

### The targeted deletion of *pflM* cures diverse *P. aeruginosa* isolates of their Pf prophages

We hypothesized that deleting *pflM* would provide a convenient way to cure *P. aeruginosa* clinical isolates of their Pf prophages. To test this hypothesis, we deleted *pflM* from the Pf prophages in cystic fibrosis isolate LESB58, two cystic fibrosis isolates from the Stanford Cystic Fibrosis Center (CPA0053 and CPA0087), and the multidrug-resistant urine isolate DDRC3 (**Table 1**).

**Table 1.** *P. aeruginosa* isolates and Pf prophage characteristics.

| Strain | Accession | Source | Pf name / lineage | Pf integration site | Pf prophage length (kb) |
| --- | --- | --- | --- | --- | --- |
| PAO1 | GCF_000006765.1 | Lab strain | Pf4 / I | tRNA-Gly | 12.4 |
|  |  |  | Pf6 / I | tRNA-Met | 12.1 |
| LESB58 | FM209186.1 | CF isolate, Liverpool, United Kingdom | Pf-LESB58 / II | Direct repeat | 10.5 |
| CPA0053 | CP137561 | CF isolate, Stanford, CA, USA | Pf-CPA0053/ II | Direct repeat | 10.4 |
| CPA0087 | CP137562 | CF isolate, Stanford, CA, USA | Pf-CPA0087/ II | tRNA-Gly | 11.1 |
| DDRC3 | CP137563 | Urine isolate, Trivandrum, Kerala, India | Pf-DDRC3/ II | tRNA-Gly | 15.5 |

Pf prophage loss was confirmed by long-read whole-genome sequencing. Targeting *pflM* successfully cured all the above clinical *P. aeruginosa* isolates of their Pf prophages (**Fig. 2A–D**). Of the Pf prophages we deleted, four were integrated into tRNA genes (three in tRNA-Gly and one in tRNA-Met) and two were integrated into direct repeats (**Table 1**). Further, Pf prophages fall into two main lineages (I and II, **Table 1**) (4) and we were successful in deleting representatives from each lineage. These observations indicate integration site nor lineage have no influence on *pflM*-mediated Pf prophage deletion.

**Figure 2:**
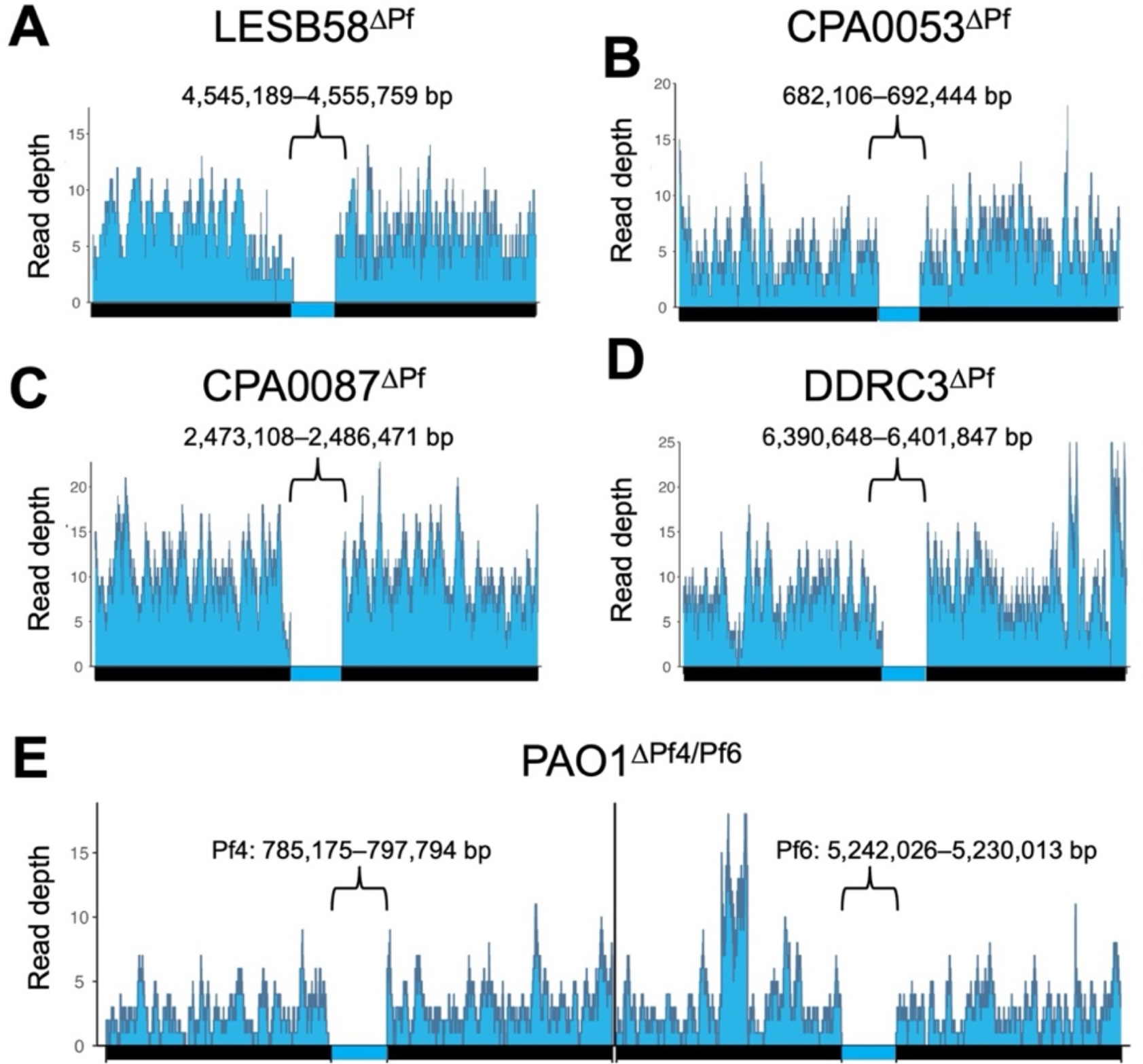
Targeted deletion of *pflM* cures diverse *P. aeruginosa* isolates of their Pf prophage infections. **(A-E)** Long-read whole genome sequencing was used to confirm the successful deletion of the indicated Pf prophages. Reads were aligned to 50kb sequences flanking the Pf prophage insertion sites in the parental chromosome. The genomic coordinates for each Pf prophage are shown above each bracket.

**Figure S1.**
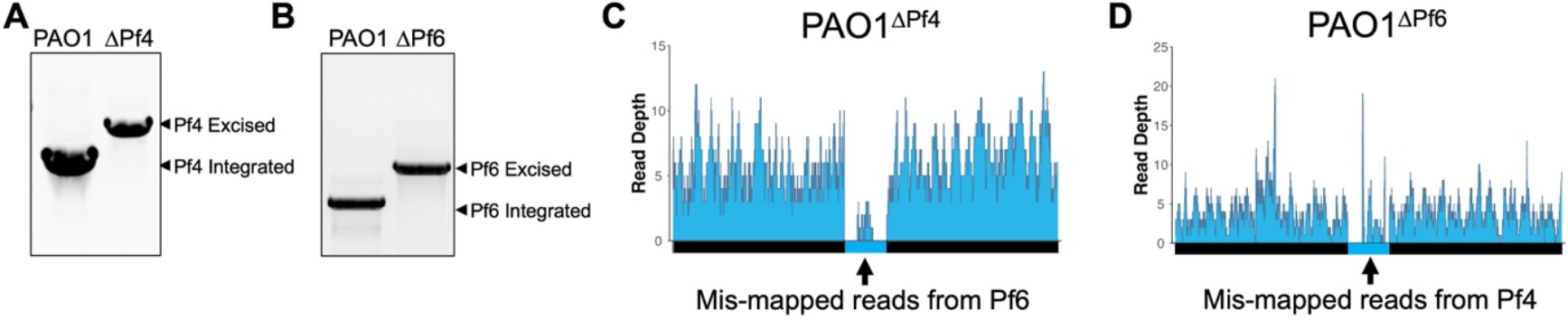
Confirmation of PAO1^ΔPf4^ and PAO1^ΔPf6^ single prophage mutants. **(A and B)** A multiplex PCR assay was used to confirm excision of (A) the Pf4 prophage or (B) the Pf6 prophage from the PAO1 chromosome. **(C and D)** Pf prophage mutants were sequenced by long-read sequencing. Arrows indicate mis-mapped reads from Pf6 to Pf4 in (C) and Pf4 to Pf6 in (D).

Many *P. aeruginosa* strains are infected by one or more Pf prophages (3). For example, some *P. aeruginosa* PAO1 sub-isolates are infected by Pf4 and Pf6 (19). Deleting *pflM* from Pf4 results in the loss of the Pf4 prophage, as does deleting *pflM* from Pf6 (**Fig** .**2A, B, Fig. S1**). Furthermore, we were able to delete Pf6 from ΔPf4, producing a PAO1^ΔPf4/Pf6^ double mutant (**Fig 2E**). This observation indicates that *pflM* from one Pf prophage is specific to that prophage and does not compensate for the loss of *pflM* from another Pf prophage residing in the same host.

PflM specificity may be explained by the diversity in the operon encoding *pflM*. In Pf4, *pflM* is truncated by a 5’ insertion of *PA0717* (4) (**Fig. 3**). The truncated *pflM* allele in Pf4, Pf LESB58, and Pf CPA0053 contain a predicted DUF5447 domain (pfam17525) whereas the *pflM* allele in other Pf strains, such as Pf6, Pf CPA0053, and Pf DDRC3, contain an additional Arc/Mnt domain (**Fig. 3**). Arc/Mnt proteins encoded by *Salmonella* phage P22 govern lysis-lysogeny decisions by binding phage operator sequences (20, 21), suggesting PflM may regulate Pf lysis-lysogeny decisions by a similar mechanism.

**Figure 3.**
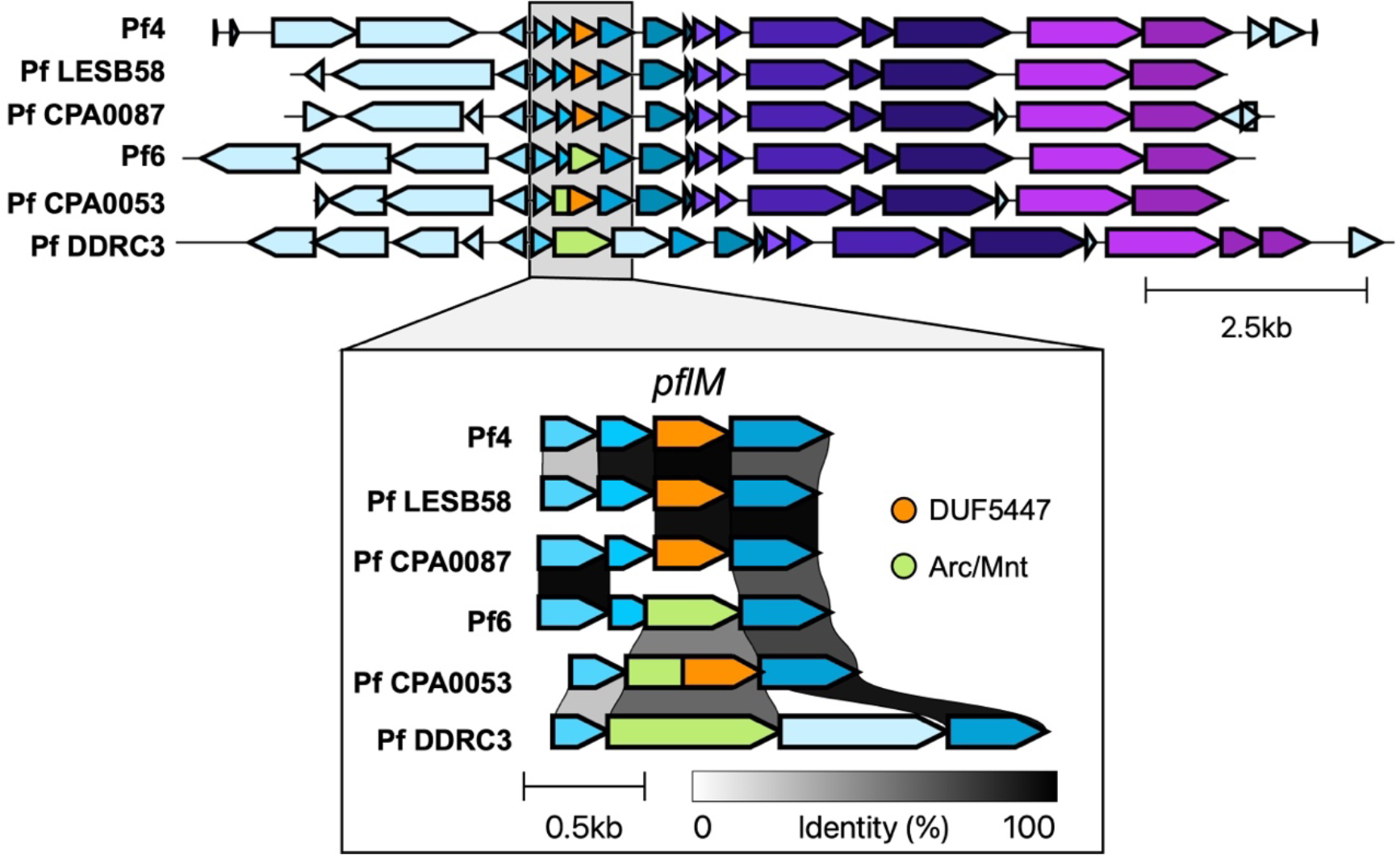
Variation in the operon encoding *pflM* in Pf prophages.

### Pf phages differentially modulate host quorum sensing

Pf4 is known to suppress PQS and Rhl quorum sensing in *P. aeruginosa* PAO1 (12, 13, 22). We hypothesized that Pf phages would likewise modulate quorum sensing in the Pf deletion strains we constructed here. To test this, we used fluorescent transcriptional reporters (12) to measure Las (*rsaL*), Rhl (*rhlA*), and PQS (*pqsA*) transcriptional activity in parental strains and Pf mutants.

We find that differences in quorum sensing activity vary by Pf strain and host. In PAO1^ΔPf4^, Las signaling is not significantly affected, Rhl transcription is downregulated, and PQS is upregulated (**Fig. 4A**). Pf6 differentially affects host quorum sensing—Las, Rhl, and PQS signaling are all upregulated in PAO1^ΔPf6^ compared to the parental strain (**Fig. 4A**). Deleting both Pf4 and Pf6 had no significant impact on Las or Rhl signaling, but PQS signaling was significantly upregulated in PAO1^ΔPf4/Pf6^ compared to the parental strain (**Fig. 4A**).

**Figure 4:**
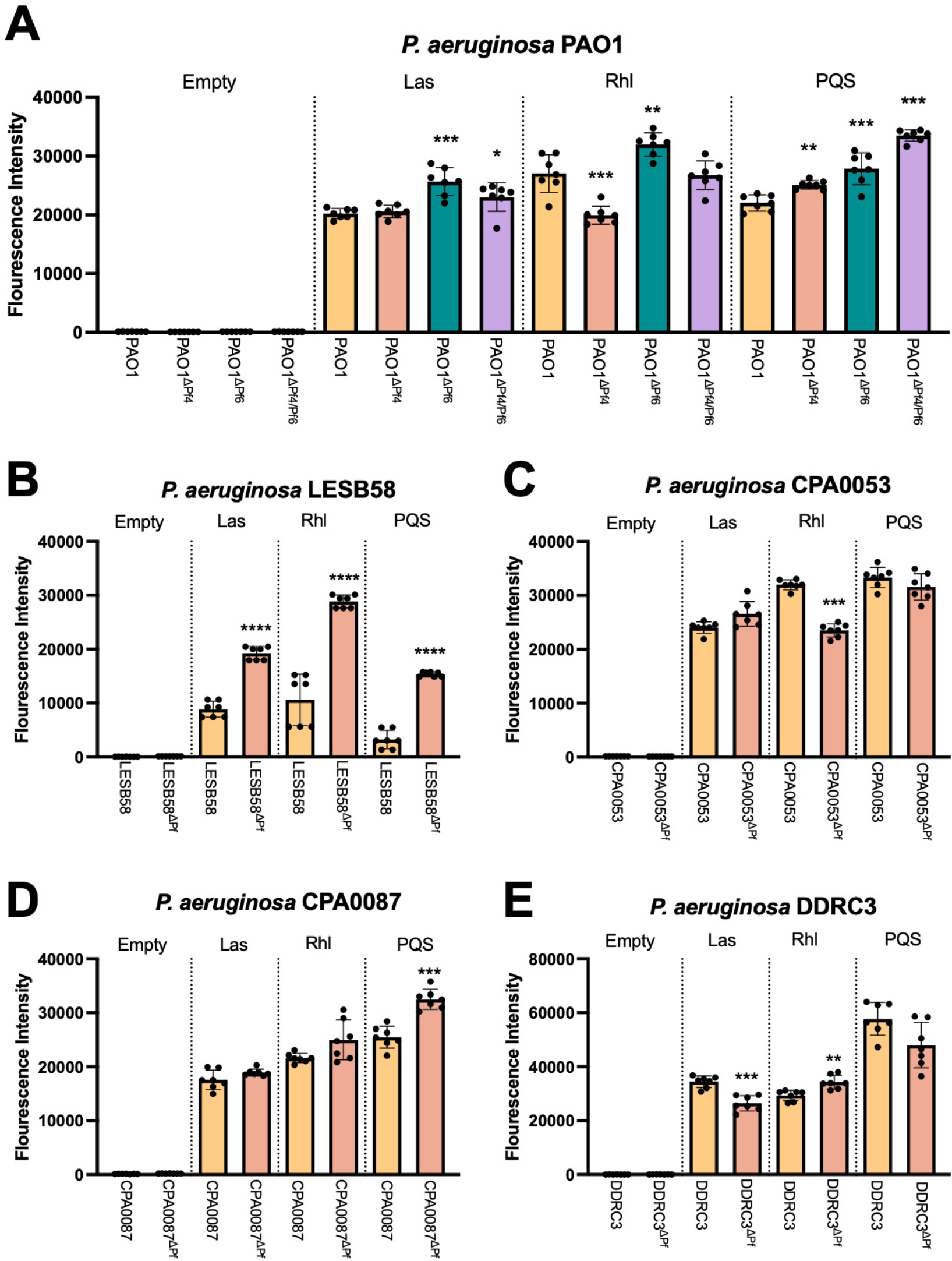
Pf phage differentially modulate *P. aeruginosa* quorum sensing. **(A-E)** GFP fluorescence from the transcriptional reporters P_*rsaLI*_*-gfp* (Las), P_*rhlA*_*-gfp* (Rhl), or P_*pqsA*_*-gfp* (PQS) was measured in the indicated strains after 18 hours of growth. GFP fluorescence intensity was normalized to cell growth (OD_600_). Data are the mean ±SEM of seven replicates. *P<0.05, **P<0.01, ***P<0.001, ****P<0.0001, Student’s *t*-test comparing ΔPf strains to the wild-type parent.

PQS transcriptional activity is also significantly (P<0.0001) upregulated in LESB58^ΔPf^ as are Las and Rhl (**Fig 4B**). In strain CPA0053, Las and PQS signaling is not significantly affected while Rhl transcription is reduced when the Pf prophage is deleted (**Fig. 4C**). PQS is activated in CPA0087^ΔPf^ while Las and Rhl signaling is not significantly affected (**Fig. 4D**). Finally, in DDRC3^ΔPf^, Las is downregulated, Rhl signaling is upregulated, and PQS signaling is not significantly affected compared to the parental DDRC3 strain but is trending downward (**Fig. 4E**). Taken together, these data indicate that Pf phages have diverse and complex relationships with host quorum sensing systems that vary significantly by strain.

### Pf phages have contrasting impacts on *P. aeruginosa* biofilm formation

Pf4 is known to promote *P. aeruginosa* PAO1 biofilm assembly and function (3, 5, 7, 11, 14, 23, 24). To test if other strains of Pf affect biofilm formation, we used the crystal violet biofilm assay (25) to measure biofilm formation of lysogenized *P. aeruginosa* isolates compared to the Pf prophage deletion mutants. In PAO1, deletion of either Pf4 or Pf6 significantly (P<0.001) reduce biofilm formation by 1.79- and 2.33-fold, respectively, while deletion of both Pf4 and Pf6 reduces biofilm formation by 7.14-fold (**Fig. 5A**). This result indicates both Pf4 and Pf6 contribute to PAO1 biofilm formation, which is consistent with prior observations (5, 7, 11, 14, 23, 24). The clinical isolates in general did not form as robust biofilms as the PAO1 laboratory strain under the *in vitro* conditions tested. Even so, deleting the Pf prophage from strains CPA0053 and DDRC3 modestly but significantly (P<0.05) reduced biofilm formation (**Fig. 5B and C**). In contrast, biofilm formation was significantly (P<0.01) increased in strains LESB58^ΔPf^ and CPA0087^ΔPf^ compared to the parental strains (**Fig. 5D and E**). The variation in biofilm formation phenotypes is perhaps not surprising given the variation in quorum sensing regulation between Pf lysogens and their corresponding Pf prophage mutants (**Fig 4**).

**Figure 5:**
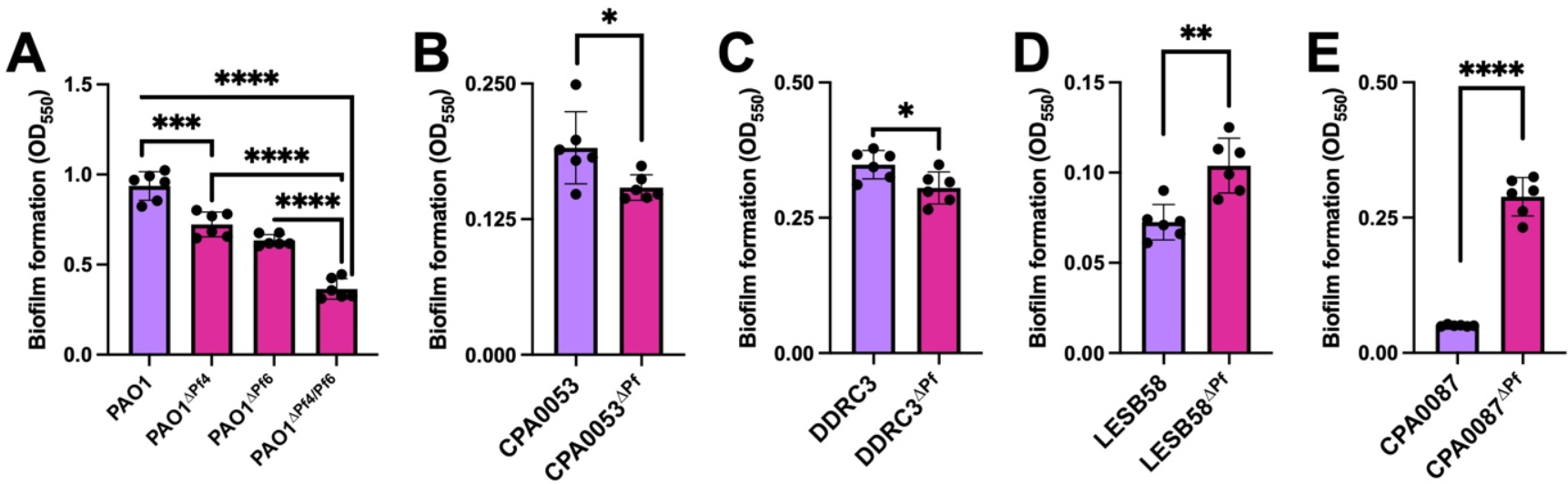
Pf prophage deletion has significant but variable effects on *P. aeruginosa* biofilm formation. Crystal violet biofilm assays were performed to measure biofilm formation of the indicated strains after 48h incubation. Data are the mean ±SEM of six replicates. *P<0.05, **P<0.01, ***P<0.001, ****P<0.0001, Student’s *t*-test.

### Pf phages suppress *P. aeruginosa* pyocyanin production

Pyocyanin is a redox-active quorum-regulated virulence factor (26). Deleting the Pf4 prophage from PAO1 enhances pyocyanin production (27). We observed increased pyocyanin production in all ΔPf strains tested except CPA0053, which did not produce much pyocyanin under any condition tested (**Fig. 6A-E**). These results suggest that Pf prophages encode gene(s) that inhibit host pyocyanin production, which is consistent with recent work indicating that the PfsE protein encoded by Pf phages inhibits PQS signaling by binding to and inhibiting PqsA (13).

**Figure 6:**
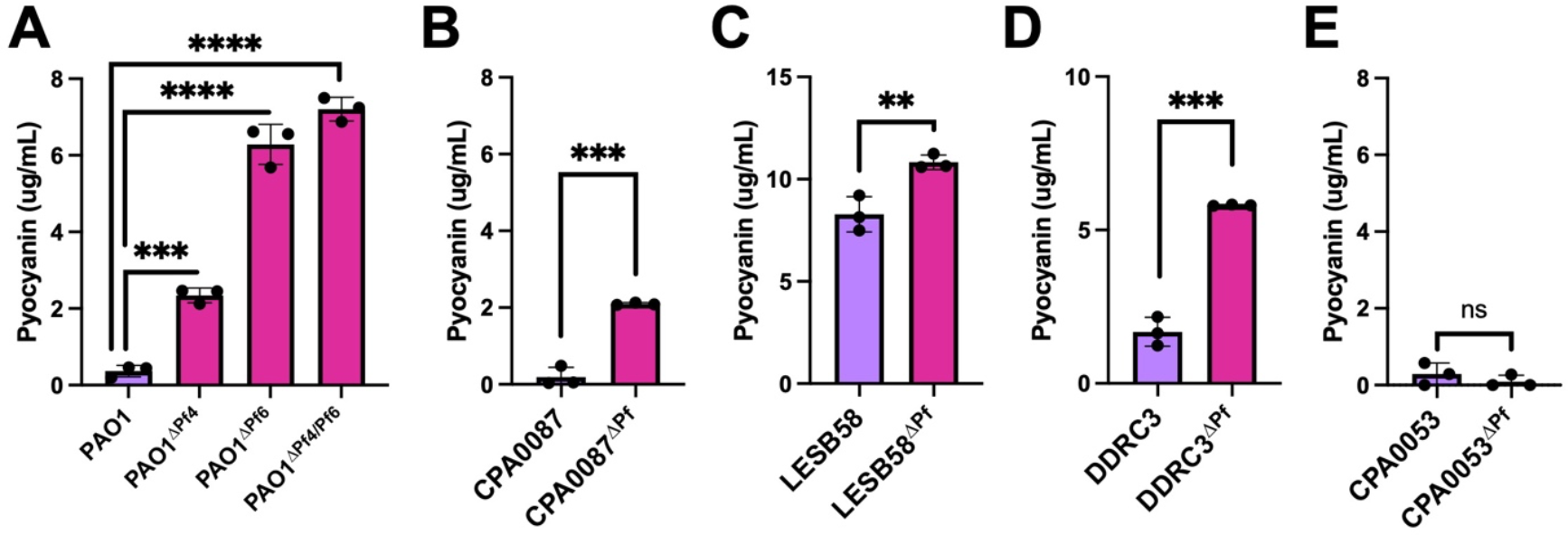
Pyocyanin production is enhanced in Pf prophage deletion strains. **(A-E)** Pyocyanin was CHCl_3_-HCl extracted from the supernatants of the indicated cultures after 18h of incubation. Pyocyanin concentration was measured (Abs 520nm). Data are the mean ±SEM of three replicates. **P<0.01, ***P<0.001, ****P<0.0001, Student’s *t*-test.

### Pf phages induce avoidance behavior in bacterivorous nematodes

In the environment, bacterivores impose high selective pressures on bacteria (28, 29). Pf4 modulation of quorum-regulated virulence factors increases *P. aeruginosa* fitness against the bacterivorous nematode *Caenorhabditis elegans* (12). We hypothesized that Pf prophages in *P. aeruginosa* clinical isolates would similarly protect *P. aeruginosa* from predation by *C. elegans*. To test this, we employed *C. elegans* avoidance assays (30-33) as a metric of bacterial fitness when confronted with nematode predation (**Fig. 7A**). *C. elegans* avoided all Pf lysogens, preferring to associate with the ΔPf strains in every case (**Fig. 7B**). Note that nematode survival was over 95% over the course of the experiment (8 hours) in all experiments (**Fig. 7B**, triangles). Collectively, our results suggest that Pf modulates *P. aeruginosa* virulence phenotypes in ways that repel *C. elegans*.

**Figure 7:**
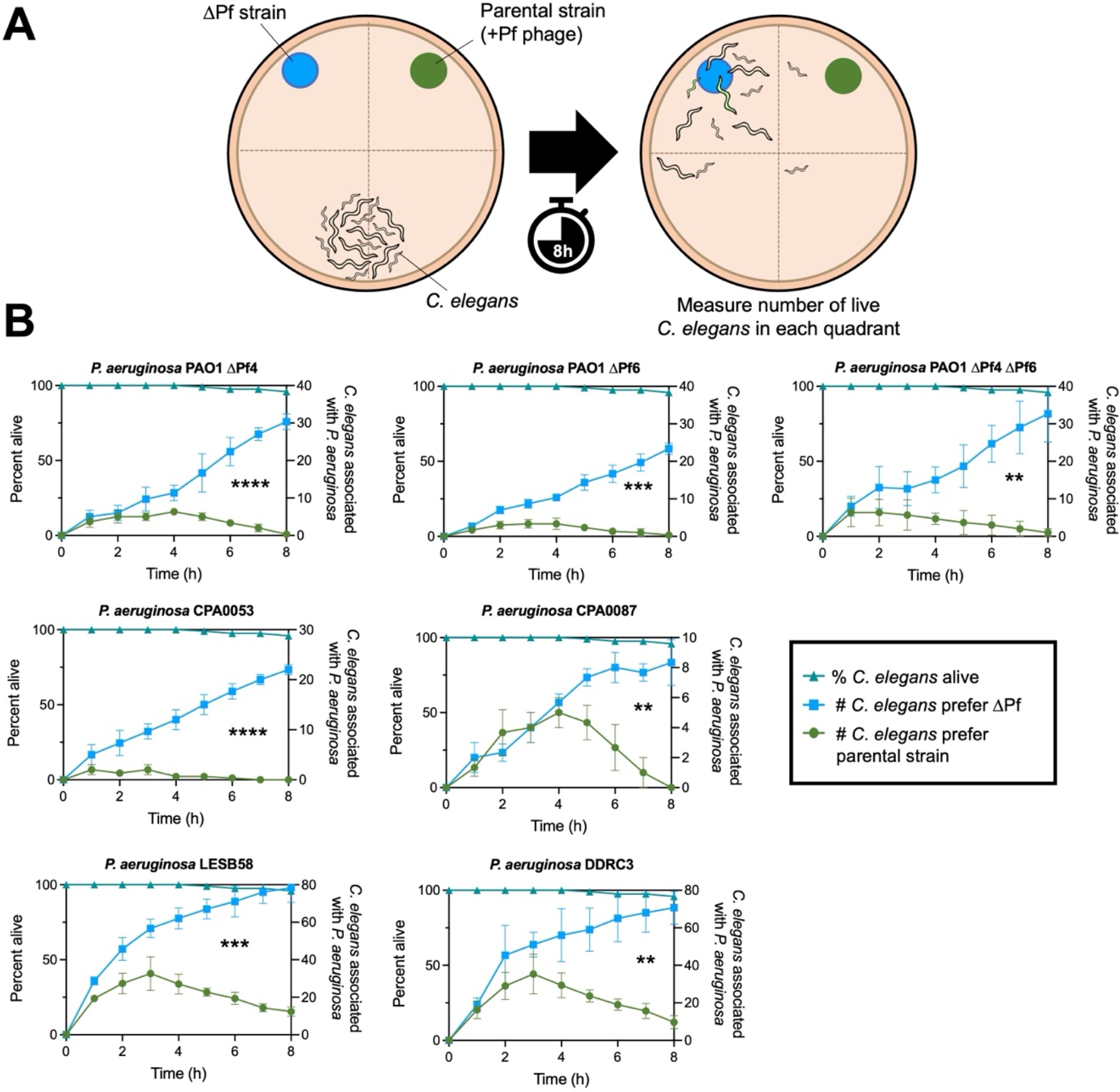
*C. elegans* actively avoids *P. aeruginosa* Pf lysogens. **(A)** Experimental design: *P. aeruginosa* and an isogenic ΔPf mutant were spotted onto NMMG plates with wild-type N2 *C. elegans* at the indicated locations. *C. elegans* localization to the indicated quadrants was measured hourly. **(B)** *C. elegans* association with *P. aeruginosa* (circles) or isogenic ΔPf mutants (squares) in the indicated strain backgrounds was measured hourly over eight hours (three experiments with N=30 per replicate [90 animals total]). P values were calculated by two-way ANOVA comparing ΔPf strains to the parental strains using the Šidák correction (95% CI threshold), **P<0.01, ***P<0.001, ****P<0.0001.

## Discussion

This study describes a convenient method to cure *P. aeruginosa* isolates of their Pf prophage infections and explores relationships between diverse Pf phages and their *P. aeruginosa* hosts. Overall, different Pf strains exhibit varying effects on host quorum sensing and biofilm formation. One commonality between all Pf strains examined is their ability to suppress pyocyanin production and repel *C. elegans* away from *P. aeruginosa*, protecting their host from predation.

Our results indicate that PflM maintains Pf in a lysogenic state and that deleting the *pflM* gene induces Pf prophage excision, but not replication. In Pf4, the site-specific tyrosine recombinase IntF4 catalyzes Pf4 prophage integration into and excision from the chromosome while the Pf4 excisionase XisF4 regulates Pf4 prophage excision by promoting interactions between IntF4 and Pf4 attachment sites as well as inducing the expression of the replication initiation protein PA0727 (4, 17). In response to stimuli such as oxidative stress (34), these coordinated events induce Pf4 prophage excision and initiate episomal replication, allowing Pf4 to complete its lifecycle.

While it is presently not known how PflM maintains Pf lysogeny, it is possible that PflM promotes the integrase activity of IntF or inhibits XisF-mediated excision, causing the Pf prophage to excise from the chromosome without concurrently inducing the replication initiation protein PA0727 (6), thus resulting in Pf prophage excision without initiating episomal Pf replication.

Our study highlights a role for Pf phages in manipulating *P. aeruginosa* quorum sensing. Pf phages have varying effects on host quorum sensing; broadly, we determine that Pf phage modulate quorum sensing activity and quorum-regulated phenotypes in all strains tested. These findings imply that different Pf strains interact with host quorum sensing networks in diverse ways, indicating a complex interplay between Pf phages and host regulatory systems.

Quorum sensing regulates *P. aeruginosa* biofilm formation and Pf4 contributes to biofilm formation in PAO1 (5, 7, 14, 18, 35). Consistently, we find that both Pf4 and Pf6 contribute to PAO1 biofilm formation. Interestingly, the impact of Pf prophage deletion on biofilm formation varies among clinical isolates, which may be related to different quorum sensing hierarchies present in clinical *P. aeruginosa* isolates (36).

Despite differences in interactions between Pf phages and host quorum sensing, deleting Pf prophages from the host chromosome enhances pyocyanin production in all strains tested except for strain CPA0053, which produces low levels of pyocyanin compared to all other strains tested. As pyocyanin is the terminal signaling molecule in *P. aeruginosa* quorum sensing networks (26), these results suggest that inhibition of pyocyanin production is the ultimate goal of Pf phages and inhibition of pyocyanin production may be beneficial to Pf phages during active replication. Pyocyanin and other redox-active phenazines are toxic to bacteria; it is possible that stress responses that are induced by pyocyanin-producing *P. aeruginosa* are detrimental to Pf replication.

We recently discovered that Pf phages encode a protein called PfsE (PA0721) that inhibits PQS signaling by binding to the anthranilate-coenzyme A ligase PqsA and that this results in enhanced Pf replication (13). It is possible that the loss of PfsE in the ΔPf strains in this study is responsible for the observed increase in pyocyanin production.

Pf lysogens induce avoidance behavior by *C. elegans*, which prefers to associate with the ΔPf strains. Strikingly, although strains lacking Pf prophages are less virulent in a nematode infection model, the reduced virulence of ΔPf strains contrasts with their high pyocyanin virulence factor production. This discrepancy may be partly explained by our prior findings that Pf4 suppresses pyocyanin and other bacterial pigment production as a means to avoid detection by innate host immune responses (12) that are regulated by the aryl hydrocarbon receptor (37, 38).

In summary, this research reveals the crucial role of the PflM gene in maintaining Pf lysogeny, demonstrates strain-specific effects on quorum sensing and biofilm formation, reveals the consistent inhibition of pyocyanin production by Pf phages, and suggests a role for Pf phages in protecting *P. aeruginosa* against nematode predation.

## Materials and Methods

### Strains, plasmids, primers, and growth conditions

Strains, plasmids, and their sources are listed in **Table 2**. Unless otherwise indicated, bacteria were grown in lysogeny broth (LB) at 37 °C with 230 rpm shaking and supplemented with gentamicin (Sigma) where appropriate, at either 10 or 30 μg ml^−1^.

**Table 2.**
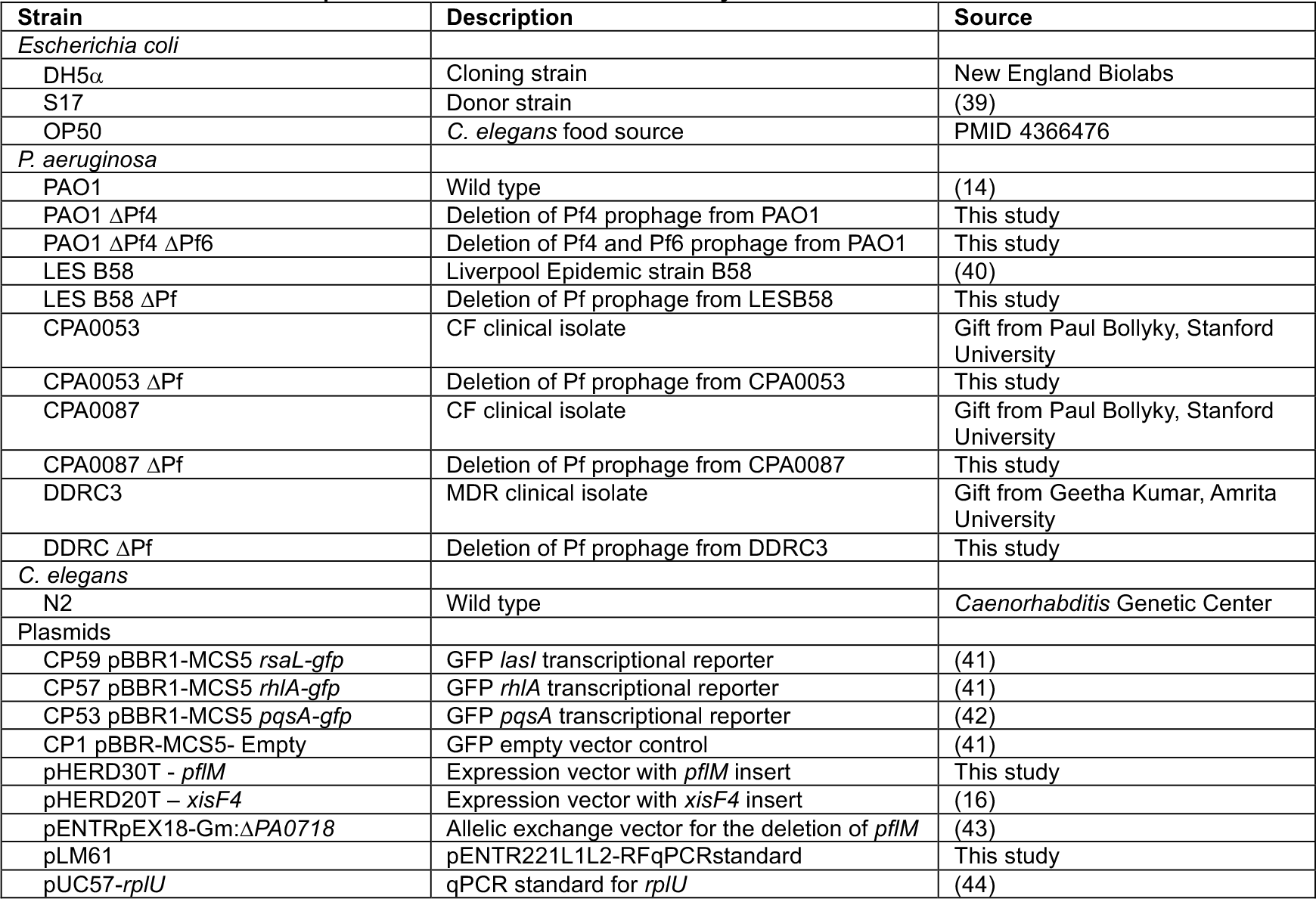
Strains and plasmids used in this study.

### Construction of Deletion Mutants

We used allelic exchange to delete alleles from *P. aeruginosa* (43). Briefly, to delete *pflM* (*PA0718)* upstream and downstream homologous sequences (∼500bp) were amplified through PCR from PAO1 genomic DNA using the UP and DOWN primers listed in **Table 3**. These amplicons were then ligated through splicing-by-overlap extension (SOE)-PCR to construct a contiguous deletion allele. This amplicon was then run on a 0.5% agarose gel, gel extracted (*New England Biolabs #T3010L*), and cloned (Gateway, Invitrogen) into a pENTRpEX18-Gm backbone to produce the deletion construct. The deletion construct was then transformed into DH5α, mini-prepped (*New England Biolabs #T1010L*), and sequenced (Plasmidsaurus.com). Sequencing-confirmed vectors were then transformed into *E. coli* S17 Donor cells for biparental mating with the recipient *P. aeruginosa* strain. Single crossovers were isolated on VBMM agar supplemented with 30 μg/mL gentamicin followed by selection of double crossovers on no salt sucrose. The final obtained mutants were confirmed by excision assay (see below), Sanger sequencing of excision assay products, and whole genome sequencing.

**Table 3.**
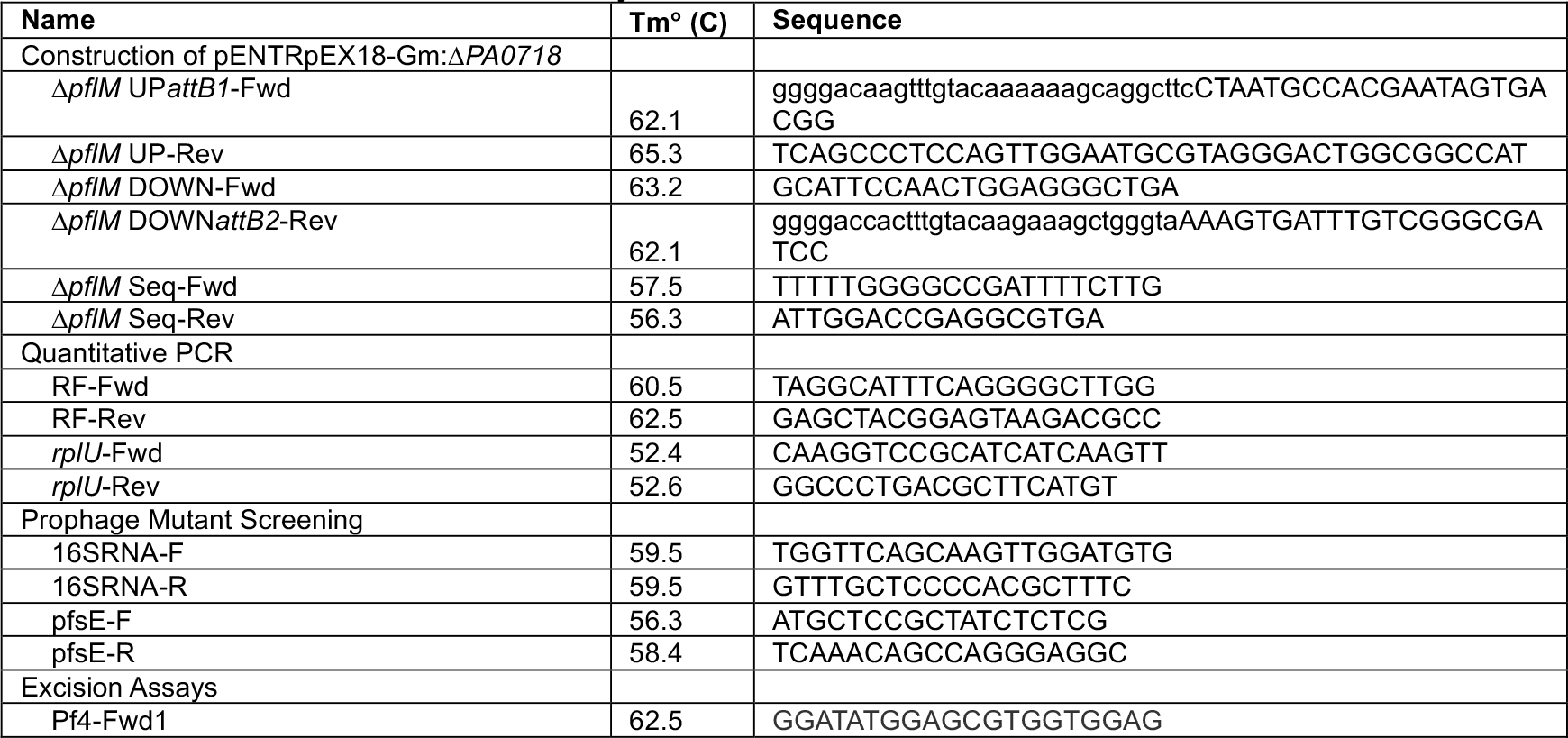
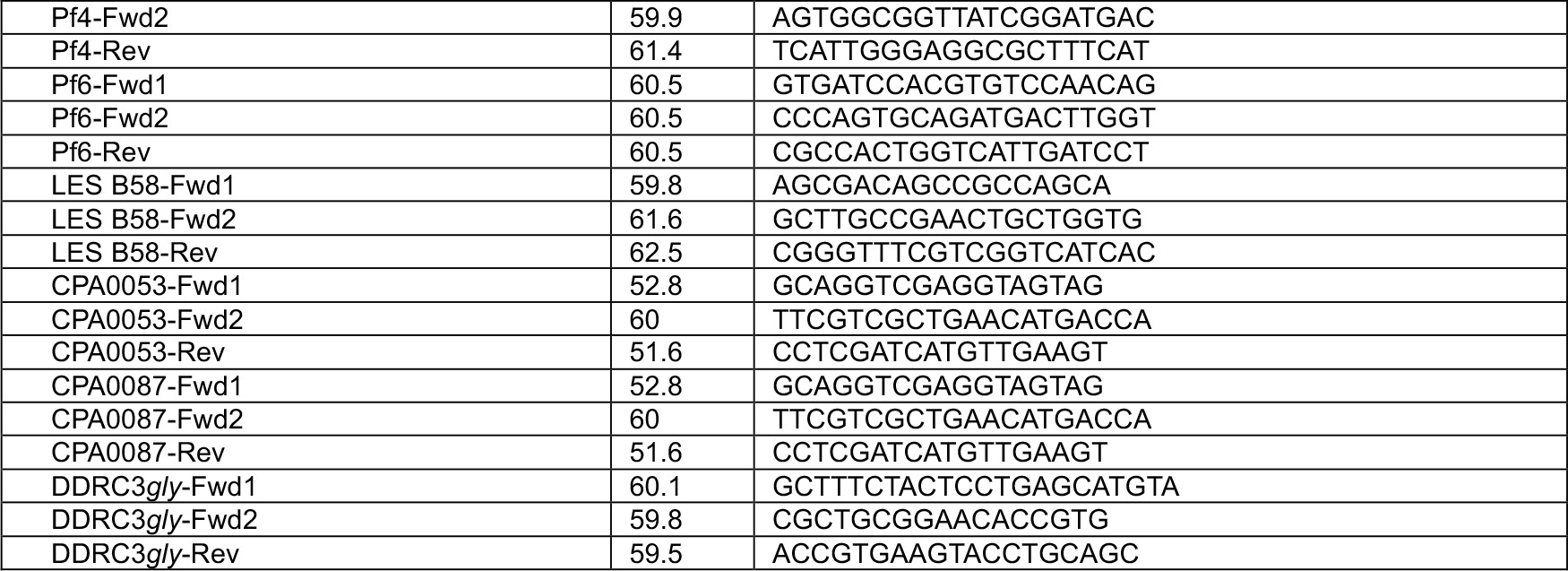
Primers used in this study.

### Excision Assays

Excision assays were designed as described previously (45). Briefly, a multiplex PCR assay was designed to produce amplicons of distinct sizes if the Pf prophage was integrated (primers Fwd_1 and Rev produce a smaller band) or excised (primers Fwd_2 and Rev produce a larger band) using Phusion Plus PCR Mastermix (Thermo Scientific # F631L). Primers were used at a final concentration of 0.5μM and are listed in (**Table 3**).

### Plaque assays

Plaque assays were performed using ΔPf4 as the indicator strain grown on LB plates. Phage in filtered supernatants were serially diluted 10x in PBS and spotted onto lawns of PAO1^ΔPf4^. Plaques were imaged after 18h of growth at 37 °C. PFUs/mL were then calculated.

### Quantitative PCR (qPCR)

Cultures were grown overnight in LB broth with shaking at 37°C. Following 18h incubation, cultures were pelleted at 16,000xg for 5 minutes, washed 3x in 1X PBS, and treated with DNase at a final concentration of 0.1 mg/mL. qPCR was performed using SsoAdvanced Universal SYBR Green Supermix (BioRad #1725270) on the BioRad CFX Duet. For the standard curves, the sequence targeted by the primers were inserted into vectors pLM61 and pUC57-*rplU*, respectively, and 10-fold serial dilutions of the standard were used in the qPCR reactions with the appropriate primers (**Table 3**) to construct standard curves. Normalization to chromosomal copy number was performed as previously described (44) using 50S ribosomal protein gene *rpIU*.

### Pyocyanin extraction and measurement

Pyocyanin was measured as previously described (46, 47). Briefly, 18-hour cultures were treated with chloroform at 50% vol/vol. Samples were vortexed vigorously and the organic phase separated by centrifuging samples at 6,000xg for 5 minutes. The chloroform layer was removed to a fresh tube and 20% the volume of 0.1 N HCl was added and the mixture vortexed vigorously. Once separated, the aqueous fraction was aliquoted to a 96-well plate and absorbance measured at 520 nm. The concentration of pyocyanin, expressed as μg/ml, was obtained by multiplying the OD_520_ nm by 17.072, as described previously (47).

### Quorum sensing reporters

Competent *P. aeruginosa* cells were prepared by washing overnight cultures in 300 mM sucrose followed by transformation by electroporation (48) with the plasmids CP1 PBBR-MCS5 *Empty*, CP53 PBBR1-MCS5 *pqsA-gfp*, CP57 PBBR1-MCS5 *rhlA-gfp*, CP59 PBBR1-MCS5 *rsaL-gfp* listed in (**Table 2**). Transformants were selected by plating on the appropriate antibiotic selection media. The indicated strains were grown in buffered LB containing 50 mM MOPS and 100 μg ml^−1^ gentamicin for 18 hours. Cultures were then sub-cultured 1:100 into fresh LB MOPS buffer and grown to an OD_600_ of 0.3. To measure reporter fluorescence, each strain was added to a 96-well plate containing 200 μL LB MOPS with a final bacterial density of OD_600_ 0.1 and incubated at 37 °C in a CLARIOstar BMG LABTECH plate reader. Prior to each measurement, plates were shaken at 230 rpm for a duration of two minutes. A measurement was taken every 15 minutes for both growth (OD_600_) or fluorescence (excitation at 485–15 nm and emission at 535–15 nm). End-point measurements at 18h were normalized to cell density.

### *C. elegans* Growth Conditions

Synchronized adult N2 *C. elegans* were propagated on Normal Nematode Growth Medium (NNGM) agar plates with *E. coli* OP50 as a food source.

### *C. elegans* Avoidance Assays

*C. elegans* avoidance assays were performed as previously described (32). Briefly, synchronized adult N2 worms were propagated at 24°C on 3.5 cm NNGM agar plates with *E. coli* OP50 for 48h, collected, and washed 4x to remove residual OP50. NNGM agar was spotted with 20 μL of *P. aeruginosa* (Pf lysogens and their isogenic ΔPf mutant) overnight cultures (LB broth) as shown in Fig 7A and grown for 18 hours at 37°C. Worms were plated in in triplicate and incubated at 24°C. *C. elegans* migration was monitored hourly for 8h.

### Whole Genome Sequencing & Annotation

Whole genome sequencing was performed by Plasmidsaurus. Reads were filtered using Filtlong (v0.2.1), assembled using Flye (v2.9.1) and/or Velvet (v7.0.4). Contigs polished using Medaka (v1.8.0), and annotated using Bakta (v1.6.1), Bandage (v0.8.1), and RAST (https://rast.nmpdr.org/). Domain analysis was performed using PfamScan (https://www.ebi.ac.uk/Tools/pfa/pfamscan/) against the library of Pfam HMM using an e-value cutoff of 0.01. Supporting domain models were obtained from Conserved Domain Database, and Defense Finder (49). Raw sequencing reads and assemblies for parental strains introduced in this study (**Table 2**) have been deposited as part of BioProject PRJNA1031220 in the NCBI SRA database.

### Statistical analyses

Differences between data sets were evaluated with a Student’s *t*-test (unpaired, two-tailed), or two-way ANOVA using the Šidák correction (95% CI threshold) where appropriate. P values of < 0.05 were considered statistically significant. GraphPad Prism version 9.4.1 (GraphPad Software, San Diego, CA) was used for all analyses.

## Acknowledgements

This work was supported by NIH grants R01AI138981 and P20GM103546 to PRS. DRF was supported by NSF GRFP grant 366502. The funders had no role in study design, data collection and analysis, decision to publish, or preparation of the manuscript. The authors report no conflicts of interest.

